# Potential application of phage therapy for prophylactic treatment against Pseudomonas *aeruginosa* biofilm Infections

**DOI:** 10.1101/404145

**Authors:** Aditi Gurkar, Deepak Balasubramanian, Jayasheela Manur, Kalai Mathee

**Author notes:** **Current Address** Department of Human and Molecular Genetics, Herbert Wertheim College of Medicine, Biomolecular Sciences Institute, Florida International University, Miami, United States of America (Kalai Mathee). **Present Address** Department of Medicine, University of Pittsburgh, Pittsburgh, Pennsylvania, USA. Vanderbilt University, School of Medicine, Nashville, Tennessee, USA. Herbert Wertheim College of Medicine, Florida International University, Miami, Florida, USA. **Corresponding author**, Kalai Mathee, MS, PhD.

## Abstract

The majority of the microbial activity in humans is in the form of biofilms, i.e., an exopolysaccharide-enclosed bacterial mass. Unlike planktonic cells and the cells on the surface of the biofilm, the biofilm-embedded cells are more resistant to the effects of the antibiotics and the host cellular defense mechanisms. A combination of biofilm growth and inherent resistance prevents effective antibiotic treatment of *Pseudomonas aeruginosa* infections including those in patients with cystic fibrosis. Antibiotic resistance has led to an increasing interest in alternative modalities of treatment. Thus, phages that multiply *in situ* and in the presence of susceptible hosts can be used as natural, self-limiting, and profoundly penetrating antibacterial agents. The objective of this study is to identify active phages against a collection of *P. aeruginosa* isolates (PCOR strains) including the prototype PAO1 and the isogenic constitutively alginate-producing PDO300 strains. These PCOR strains were tested against six phages (P105, P134, P140, P168, P175B, and P182). The analysis shows 69 % of the PCOR isolates are sensitive and the rest are resistant to all six phages. These phages were then tested for their ability to inhibit biofilm formation using a modified biofilm assay. The analysis demonstrated that the sensitive strains showed increased resistance, but none of the susceptible strains from the initial screening were resistant. Using the minimum biofilm eradication concentration (MBEC) assay for biofilm formation, the biofilm eradication ability of the phages was tested. The data showed that a higher volume of phage was required to eradicate preformed biofilms than the amount required to prevent colonization of planktonic cells. This data supports the idea of phage therapy more as a prophylactic treatment.

## INTRODUCTION

Phage therapy may provide a solution to the global problem of increasing resistance to conventional antibiotics (1; 2). This study analyzed the application of phage as a potential therapeutic agent against *Pseudomonas aeruginosa*, the primary cause of morbidity and mortality in cystic fibrosis (CF) patients. CF was first described in 1936 in Switzerland (3) and 1938 in the U.S.A. (4; 5). It is a fatal, inherited disease afflicting 1 in every 3500 live births in the USA (6) and occurs predominantly in Caucasians. The gene responsible for CF is located on chromosome 7 (7; 8) and encodes a protein of 1480 amino acids; it is called the cystic fibrosis transmembrane conductance regulator (CFTR) (9). CFTR is a cyclic-AMP-activated chloride ion channel in the secretory epithelia (7; 10). A defect in CFTR leads to decreased fluid secretion, and the dehydration of the epithelial surfaces leads to the pathology of the disease (11). Over-secretion of mucus into the airway leads to congestion of the respiratory tract and increased susceptibility to bronchopulmonary infection (12). In spite of extensive research, CF patients continue to suffer from these chronic diseases, which are the leading cause of their mortality (13). The median survival for these patients in the USA is 29.6 years (6). Further research may not only improve the quality of their lives but increase their median survival age (6).

CF patients are primarily infected with respiratory viruses, especially respiratory syncytial viruses, leading to acute pulmonary disease soon after birth. Respiratory syncytial virus infection decreases the pulmonary function in these patients by 30 % in approximately one month (14; 15; 16). The viral infection is followed by secondary colonization and infection by bacteria such as *Staphylococcus aureus, Haemophilus influenzae*, and *P. aeruginosa* (17). In recent times, cross infection by various pathogens such as *Burkholderia cenocepacia* complex and *Pandoraea* spp. has been of significant concern in the later stage of CF disease (18; 19). *S. aureus* and *H. influenzae* can efficiently be eradicated using oral antibiotics (20; 21), but the *P. aeruginosa* infection, which occurs in 60 to 90 % of patients with CF, is never eradicated despite intensive anti-pseudomonal treatment (17).

*P. aeruginosa* is a Gram-negative bacterium that is ubiquitous; it infects nearly every human tissue and is one of the most common causes of nosocomial pneumonia, urinary tract infections and wound sepsis (22). It is an opportunistic pathogen that affects immunocompromised patients such as those with cancer, HIV, and burns (22). It is also a leading pathogen responsible for the morbidity and mortality among patients with CF, diffused pan bronchitis, and chronic obstructive pulmonary disease. The initial and intermittent colonization of CF lungs by non-mucoid *P. aeruginosa* can be eradicated by early, aggressive antibiotic therapy. However, such treatment generally fails in later stages when the colony morphology of bacteria, isolated from sputum samples, becomes mucoid (23). The mucoid phenotype is due to overproduction of a capsule-like polysaccharide called alginate (24), and this energy-consuming production of alginate may be necessary in the formation of biofilms as this helps them to adhere to surfaces (25; 26).

Biofilms are matrix-enclosed organized microbial communities, adherent to each other and to surfaces or interfaces (27); biofilm growth is now known to be the natural mode of microbial growth. The formation of a biofilm is defined to be a developmental process, consisting of five stages: (1) attachment; (2) monolayer formation; (3) microcolony formation (4) biofilm maturation, and (5) release of planktonic cells (26; 28; 29). The mature biofilms release planktonic bacteria starting the whole process again (30). Mature biofilms are composed of cells and matrix material located in matrix-enclosed ‘towers’ and ‘mushrooms’ (28). This mode of growth produces a barrier to penetration of antimicrobial agents through the matrix, is responsible for the altered growth rate of these microbial communities, and other physiological and morphological changes that appear to favor their survival (28).

The current initial treatment for patients with acute CF infections comprises of a combination of antibiotic treatment with ciprofloxacin and inhalation of colistin for about three weeks (31; 32). The treatment for chronic infection is much more controversial, and a combination of antibiotics, including ciprofloxacin, imipenem, tobramycin, and aztreonam, is used (33). However, several drawbacks have been observed with this mode of treatment. Patients seem to develop allergies to ß-lactam antibiotics (34; 35; 36; 37), and the bacteria develop resistance (38; 39; 40; 41; 42).

*P. aeruginosa* exhibits intrinsic and acquired resistance to many structurally and functionally unrelated antibiotics. The biofilm mode of growth (43), low membrane permeability (44; 45), target alteration (38; 46), and extensive linkage of the outer membrane proteins (47; 48; 49; 50), are part of its intrinsic properties for resistance. Also, *P. aeruginosa* has acquired five efflux pumps that actively pump out the antibiotics (51; 52; 53; 54; 55). The failure to successfully eradicate *P. aeruginosa* has prompted researchers to consider alternative approaches.

Treatment with vaccines and adjuvants, or the treatment with Interferon-γ which is naturally produced by lymphocytes activated by specific antigens, or with the Chinese herbal medicine, *Daphne giraldii* Nitsche, decreases the inflammatory response and enhances the bacterial clearance in an animal model (56; 57; 58; 59). The Chinese herbal supplement ginseng also seems to be a promising alternative measure for the treatment of chronic *P. aeruginosa* lung infections in CF patients (60; 61; 62). Quorum-sensing inhibitors (63), herbal supplements (64; 65; 66) and honey (67; 68) have also been proposed as potential alternatives to treat *P. aeruginosa* infections.

Another natural antimicrobial agent, bacteriophages or (phages), shows a new hope to conquer the drug-resistant bacteria. Bacteriophages were first discovered by Earnest Hankin in 1896 and they were rediscovered by Felix d’Herelle in 1901, who named them so for their ability to infect bacteria (69). D’Herelle immediately focused on these viruses’ potential for treatment of bacterial diseases. This led to numerous research papers on phage therapy in the first half of the 20th century (69). However, a poor understanding of phage biology, difficulties in bacterial identification coupled with high specificity, and the discovery of broad-spectrum antibiotics caused phage therapy to quickly decline (70). But it has recently regained popularity, for treating a wide variety of diseases whose control with chemotherapeutic agents is difficult. Phages can be ‘lytic’ or ‘lysogenic.’ In the lytic cycle, a phage will convert the bacterial cell into a phage-producing factory releasing a large number of phages. In the lysogenic life cycle, there is no progeny produced; the phage DNA becomes part of the bacterial genome (71). It is preferable to use lytic phages in treatment because they quickly reproduce within and lyse the bacteria in their host range (72).

The use of phages as antimicrobial agents has some advantages over other current methods of microbial control. One significant advantage of phages is their narrow host range, which allows phage treatment to remove a problem organism without disturbing the local microflora (70). Also, unlike antibiotics, phages need to be administered for only a short duration. As long as susceptible bacteria are present, the number of phages increase as they work their way more in-depth in the biofilms, rather than decaying over time and distance, like antibiotics (73). Phage therapy is also inexpensive, and it is less time-consuming to obtain phages on resistant strains than discovering new and useful drugs. Phages can potentially be used in conjunction with antibiotics to delay resistance. Nevertheless, there can be disadvantages associated with phage treatment. The narrow host range of phages may pose a problem. For example, any treatment for the bacteria associated with chronically infected CF patients would require very specific and strongly lytic phages (74). Phages are present ubiquitously with their host bacteria. However, infections do occur, suggesting that a higher dosage of phages may be required (75; 76). The phage preparations used for treatment usually have some bacterial debris, which can be harmful because this debris could have potential endotoxins (74). Anti-phage antibodies seem to appear a few weeks after administering phages, which might pose a problem when dealing with the treatment of chronic infections (77).

Because the advantages of phage therapy are many, it has been continually practiced in few places since its rediscovery by D’Herelle. Phage therapy is practiced on a small scale in Poland and the Republic of Georgia, Tbilisi (77). The bacterial pathogens targeted in these institutes include *S. aureus, P. aeruginosa, Klebsiella pneumoniae* and *Escherichia coli* (78). Phages used seem to cure about 90 % of the cases studied. In these carefully documented studies, it was shown that few human subjects complained about gastrointestinal pains (79). Fevers have also been associated with phage treatment. Although the cause remains unclear, there is a possibility that the crude solution of phages used may still have bacterial debris (79). Researchers are reluctant to use phage treatment systemically due to the fear of septic shock (79).

Phages have been studied to control bacteria implicated in causing food contamination by *E. coli, Listeria monocytogenes*, and *Salmonella* spp. (Table 1). Also, phage therapy has been tested against *P. aeruginosa* infections using various model systems and humans (80; 81; 82). Most of these treatments have proved to be very effective (curing 90-95 %). Recent studies have looked at the use of phages to deliver genes into mammalian cells (83). Phages have also been looked at as vaccine candidates for hepatitis, human immunodeficiency virus and various other diseases (84). It has also been proposed that phage delivery during transplantation of organs, to treat associated infections may prove to be very helpful (85).

**Table 1:** Recent uses of phages against pathogenic bacteria.

| Source/Model organism | Bacteria Treated | References |
| --- | --- | --- |
| Meat | *E. coli* | (Dykes and Moorhead 2002) |
| Eggs | *Salmonella spp.* | (Goode *et al.*, 2003) |
| Cheese | *Listeria* | (Modi *et al.*, 2001) |
| Skin grafts in humans | *P. aeruginosa* | (Soothill 1994) |
| Chronic pulmonary infection in chicken | *P. aeruginosa* | (Meitert *et al.*, 1987) |
| Burn wound in humans | *P. aeruginosa* | (Ahmad 2002) |

All these studies using phages seem to be as successful in comparison with studies that use antibiotics. This suggests that the safety factor linked with the usage of phages for treatment, insufficient negative immune host response and its efficacy hold promise as a future treatment for the bacterial infection in CF patients. Phages that have been effective against *P. aeruginosa* have been identified and studied (Table 1). These studies have used a single phage against a single strain. However, the *P. aeruginosa* genome is very diverse (86) and thus any phage to be used for treatment needs to be effective against a myriad of strains. The first aim of this study was to identify phages that would work against a large number of *P. aeruginosa* mucoid and non-mucoid strains. The second aim was to identify whether phages could be used to prevent biofilm formation, i.e., to avoid the initial adhesion of the bacteria. Finally, we wanted to determine if phage therapy could be used to eradicate preformed biofilms.

## MATERIALS AND METHODS

### Bacterial strains, media, and culture

*P. aeruginosa* isolates that were spatially, geographically and environmentally distinct, named PCOR isolates (Table 2), were used for the experiments (87). The prototype PAO1 and the constitutively alginate-producing isogenic variant PDO300 were also included in all the experiments. All the strains were streaked on Luria-Bertani (LB) agar plates (10 g tryptone, 5 g yeast extract, 10 g NaCl and 15 g agar per liter). They were cultured in LB broth, (10 g tryptone, 5 g yeast extract, 5 g NaCl, per liter) at 37 °C. For antimicrobial susceptibility experiments, Cation Adjusted Muller Hinton Broth (CAMHB) (3 g beef extract, 17.5 g acid hydrolysate of Casein, 1.5 g starch per liter) (BD Biosciences, San Jose, CA) was used.

**Table 2:**
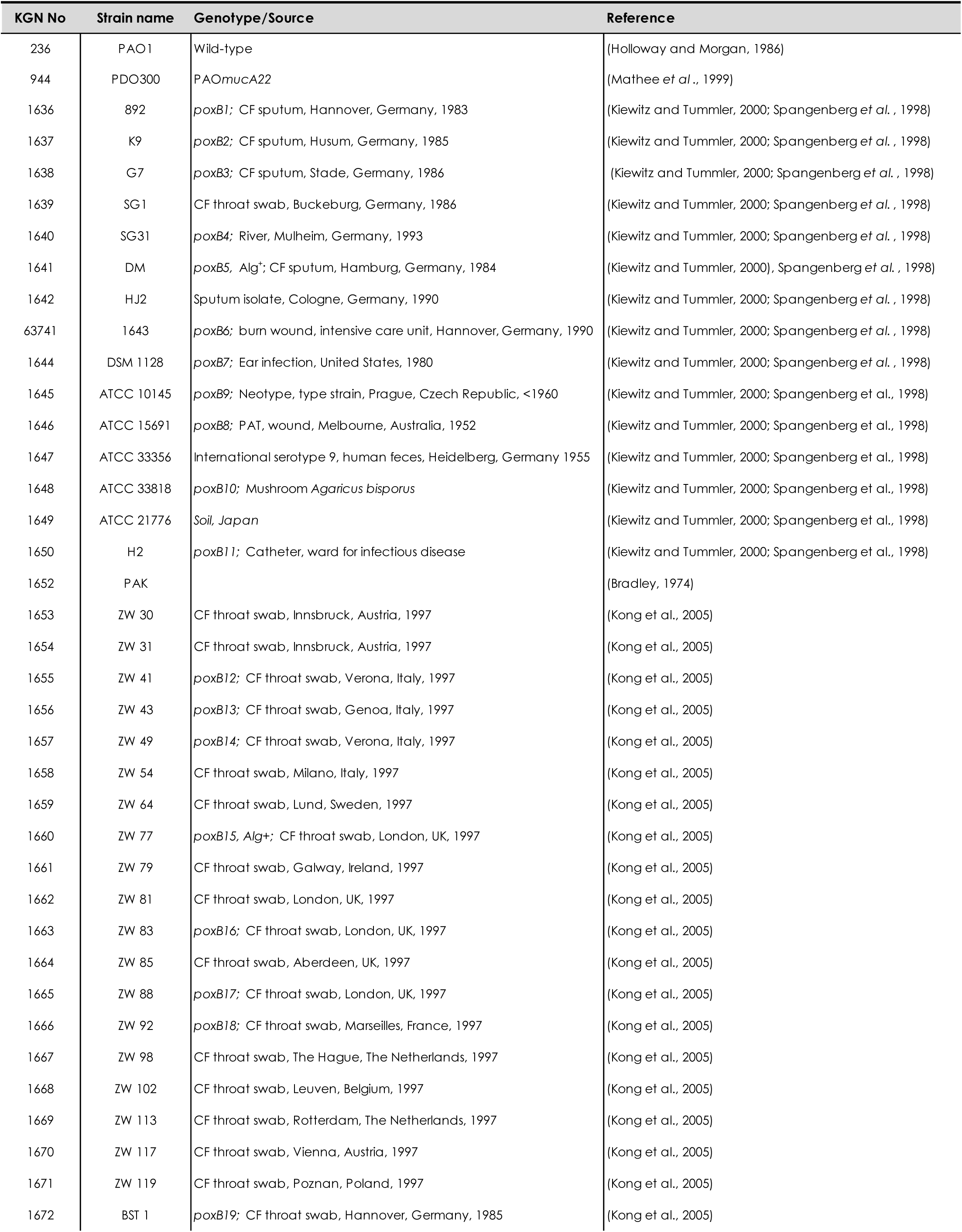
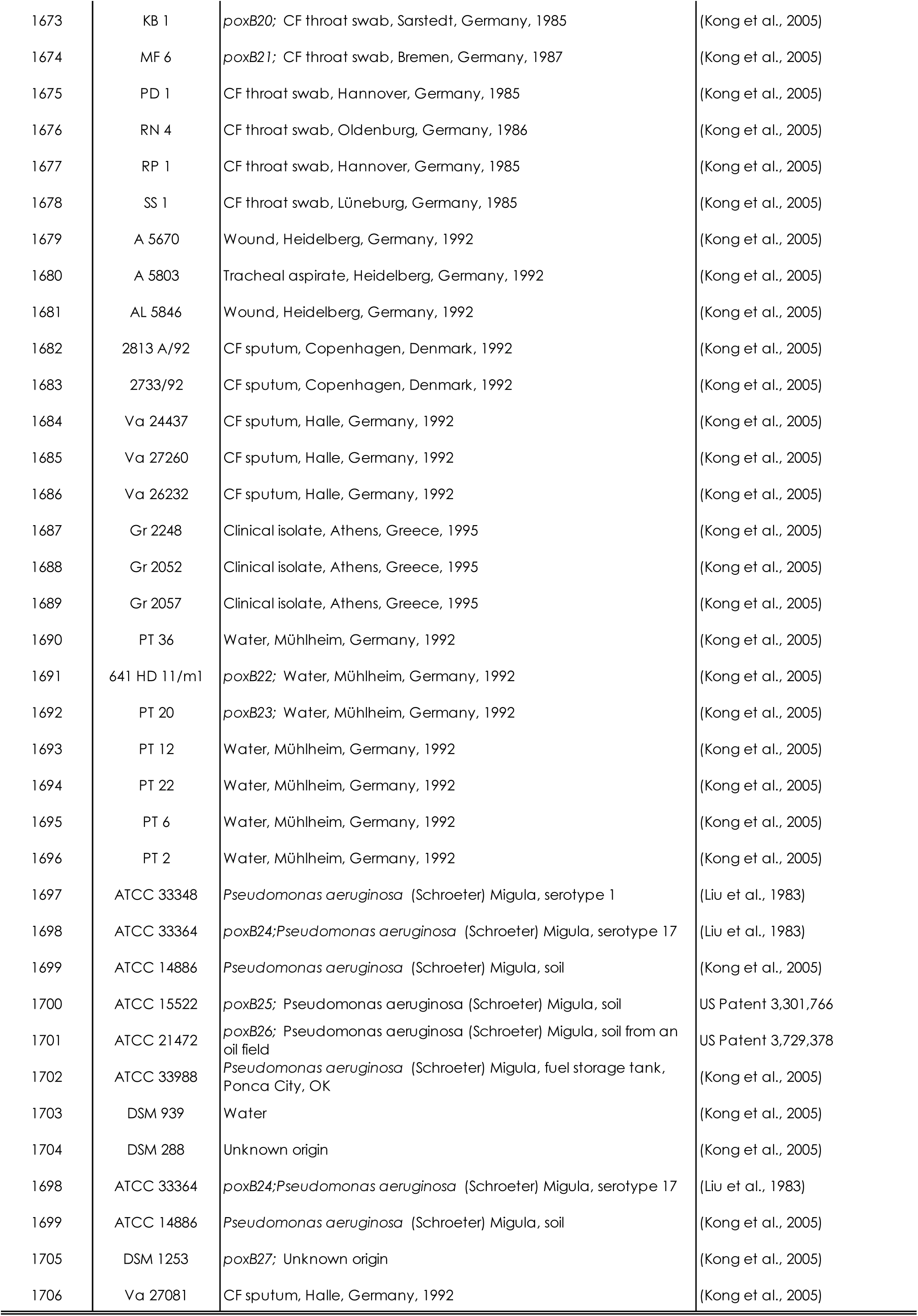
Bacterial strains, plasmids and primers used in this study.

### Phage isolation

Phages (Table 2) were isolated from the environment by GangaGen Inc, Bangalore, India. A fresh culture of the target strain (*P*. *aeruginosa*) was mixed in LB broth to which the environmental sample containing phage was added; cultures were incubated at 37 °C until complete lysis was observed. The sample was treated with 1 % chloroform and centrifuged at 16,000 *g* (Sigma Laboratory Centrifuges, 4K 15, Germany) to remove any bacterial debris.

### Phage purification

The propagating strain, the host *P. aeruginosa* PAO1, was streaked on to an LB agar plate and incubated at 37 °C for 24 h. A single colony was inoculated in 5 ml LB broth and incubated overnight at 37 °C. Fresh broth was inoculated on the following day with a 2 % inoculum; i.e., 2 ml overnight culture in a 100-ml broth. This culture was incubated in an air shaker until it reached an optical density (OD) of 0.5-0.7 as measured with a spectrophotometer (Bio-Rad SmartSpec 3000, Hercules, CA). A lawn was made by adding the bacterial strain to an LB agar plate. This LB agar plate was then allowed to dry for about 15 min. Five µL of the filtered phage was spotted onto this bacterial lawn (Figure 1). These plates were incubated for 24 h at 37 °C. A single lysed plaque from these plates was used to make pure phage stocks. Using this method, a collection consisting of over 100 phages against *P. aeruginosa* was isolated by GangaGen Inc. However, preliminary results with 11 phages suggested only six of them were required for 95 % efficacy against the PCOR isolates (data not shown). Therefore, only six phages, P105, P34, P140, P168, P175B and P182 were used for all further experiments. In order to purify these phage strains further, the prototypic strain PAO1 was used as the host cell (propagating strain) and the phage purification method was used with a slight modification as described above.

**Figure 1:**
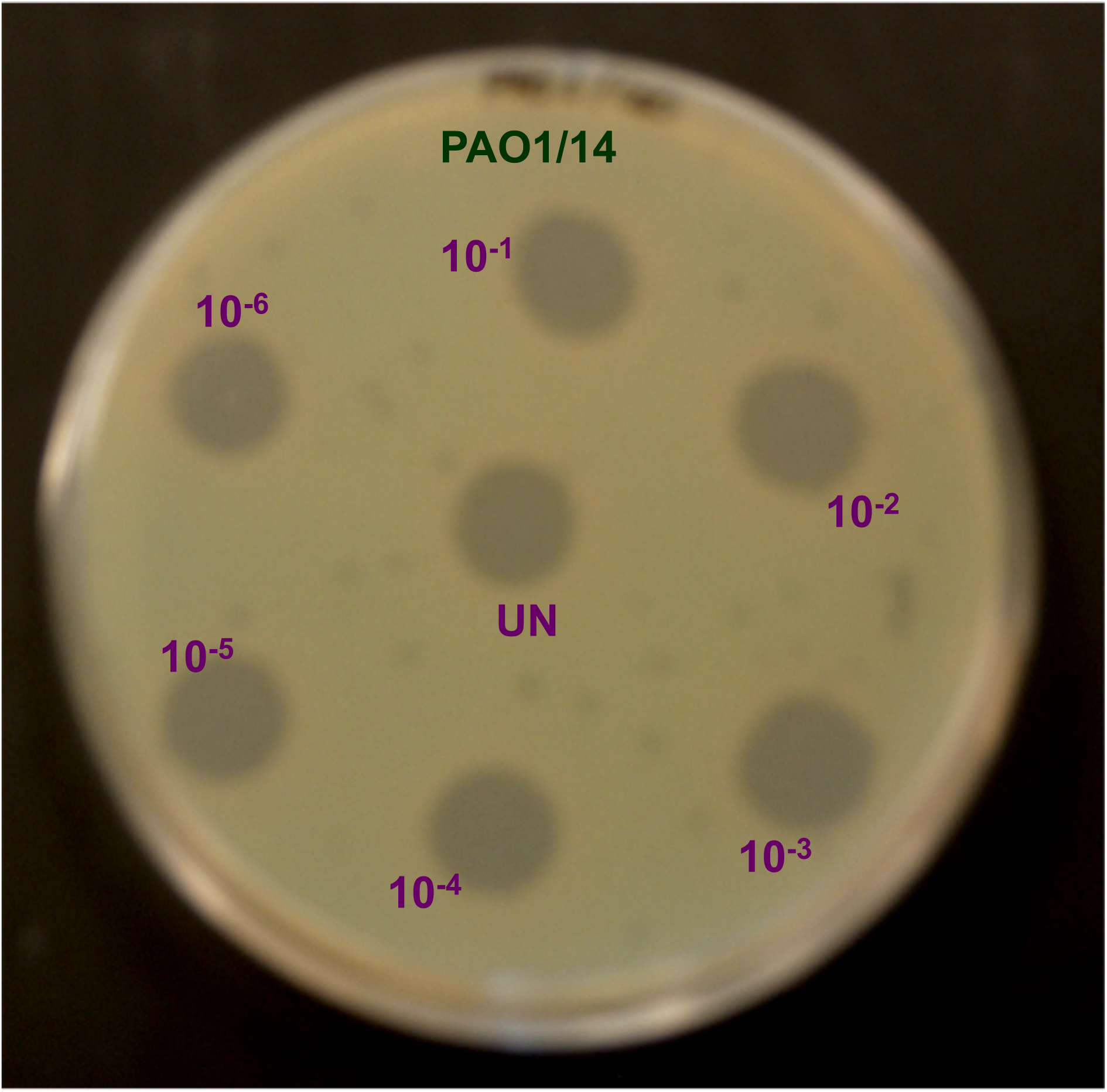
Spot titer assay. When the bacterial PAO1 lawn was seeded with phage P140 at serial dilutions, PAO1 was sensitive even at the highest dilution.

### Phage spot titer assay

A five µl serial dilution of the filtered phage was spotted on the bacterial lawn plate and incubated for 24 h (Figure 1). For further calculations of the plaque forming units (PFU/ml) and multiplicity of infection (MOI), individual plaques at suitable dilutions were counted.

### Calculation of plaque forming units (PFU/ml) and the multiplicity of infection (MOI)

Phage-infected bacterial cultures were serially diluted using LB broth. A 100-µl aliquot of an appropriate higher dilution was spread on LB agar plates, where individual plaques could be observed. The number of plaques (more than 25 and less than 300) formed on the bacterial lawn was counted. This number was used to back-calculate the approximate titer of the plaque forming units (PFU/ml).

### Phage stock preparation

PAO1 was used as the propagating strain for amplification of all phages. An overnight culture was inoculated at 2 % (V/V) in LB broth and incubated at 37 °C, shaking at 320 *g*. When the cell density reached an OD of 0.5, it was infected with the desired phage at the MOI of 0.1 and incubated at 37 °C. OD readings were taken hourly in order to follow the amount of lysis. When complete lysis (OD < 0.1) was observed, the phage solution was immediately harvested by adding 1 % of chloroform. In the absence of obvious lysis, the cultures were grown for 6 h, after which the phage-infected sample was treated with 1 % chloroform for 10 min at room temperature. This sample was centrifuged at 16,000 *g* for 20 min at 4 °C. The supernatant containing the newly-replicated phage was transferred to a new tube. A spot titer was performed to determine the new titer or PFU/ ml for each phage.

### Phage overlay

A 2 % fresh inoculum of the bacterial target strain was incubated at 37 °C and grown until it attained an OD of 0.5. The phage dilution to be tested was selected using the spot titer assay. A 100 µl aliquot of the diluted phage was added to 100 µl of the log-phase culture of the target strain and incubated for 5 min at 4°C to ensure adsorption. After which, 3 ml of soft agar was added to the adsorption mix and poured on the LB agar lawn. After an incubation period of 16 h, the numbers of plaques were counted and the PFU/ml was determined.

### Screening of PCOR isolates

Planktonic cells of 67 PCOR isolates (Table 2) that were spatially, geographically and environmentally distinct were used for this experiment. The PCOR isolates consisted of 34 CF isolates, 14 other non-CF clinical isolates and 12 strains from the environment. Along with these the prototypic nonmucoid and their isogenic mucoid variant strains, PAO1 and PDO300 were tested against the six tested phages, P105, P134, P140, P168, P175B, and P182 (Table 2). Overnight cultures of all 69 strains were inoculated in fresh LB broth until an OD 0.5 was obtained. A lawn culture of the bacterial strain to be tested was grown on LB agar. The phage stock was serially diluted using LB broth, and a 5 µl of the serial dilutions, ranging from 10^−1^ to 10^−5^, was spotted on the prepared lawn culture (Figure 1). The sensitivity of the strain to the phages was determined as follows:

- ***Sensitive***: Clear lysis at the highest dilution; the strain was sensitive at a low titer, i.e. sensitive even at a very high dilution of phage.
- ***Intermediate Sensitivity***: Lysis at a dilution of 10^−4^; the strain was sensitive at an intermediate phage titer.
- ***Low Sensitivity***: Lysis at 10^−2^; the strain was sensitive at a high phage titer.
- ***Resistant***: No lysis or lysis only at undiluted concentration; the strain was resistant.

### Biofilm inhibition assay (BIA)

Biofilm formation is the natural mode of growth for bacteria. Biofilms consists of sessile bacteria firmly attached to surfaces and each other via exopolysaccharide production conferring additional resistance to antibiotics (88). In order to identify phages that would inhibit *P. aeruginosa* biofilm formation, we tested the ability of the six isolated phages to inhibit biofilm initiation using the Biofilm Inhibition Assay (BIA) (Figure 2) (89). The phage-sensitive PCOR isolates (46/69) were examined for biofilm inhibition. The strain to be tested was diluted 1:100, and 100 µl of the diluted bacteria were inoculated into each well of a 96-well polyvinyl chloride (PVC) plate (Falcon, BD Biosciences, San Jose, CA). In addition to the bacteria, phages at increasing MOIs ranging from 0.1, 1.0, 2.5 and 5.0, were initially added to the plate. If inhibition of bacteria was not observed at an MOI of 5.0, the MOIs were further increased to 10, 20, 40 and 80. A negative control with no phage was also added to the plate. These plates were incubated at 30°C for 10 h. All samples were run in duplicate. After the incubation period, 25 µl of crystal violet (CV) was added to each well, and the plate was incubated at room temperature for 15 min. Crystal violet was absorbed by the cells adhered to the wells, indicating the formation of a biofilm (Figure 3). After 15 min, the plates were rinsed with water repeatedly to release unbound planktonic cells.

**Figure 2:**
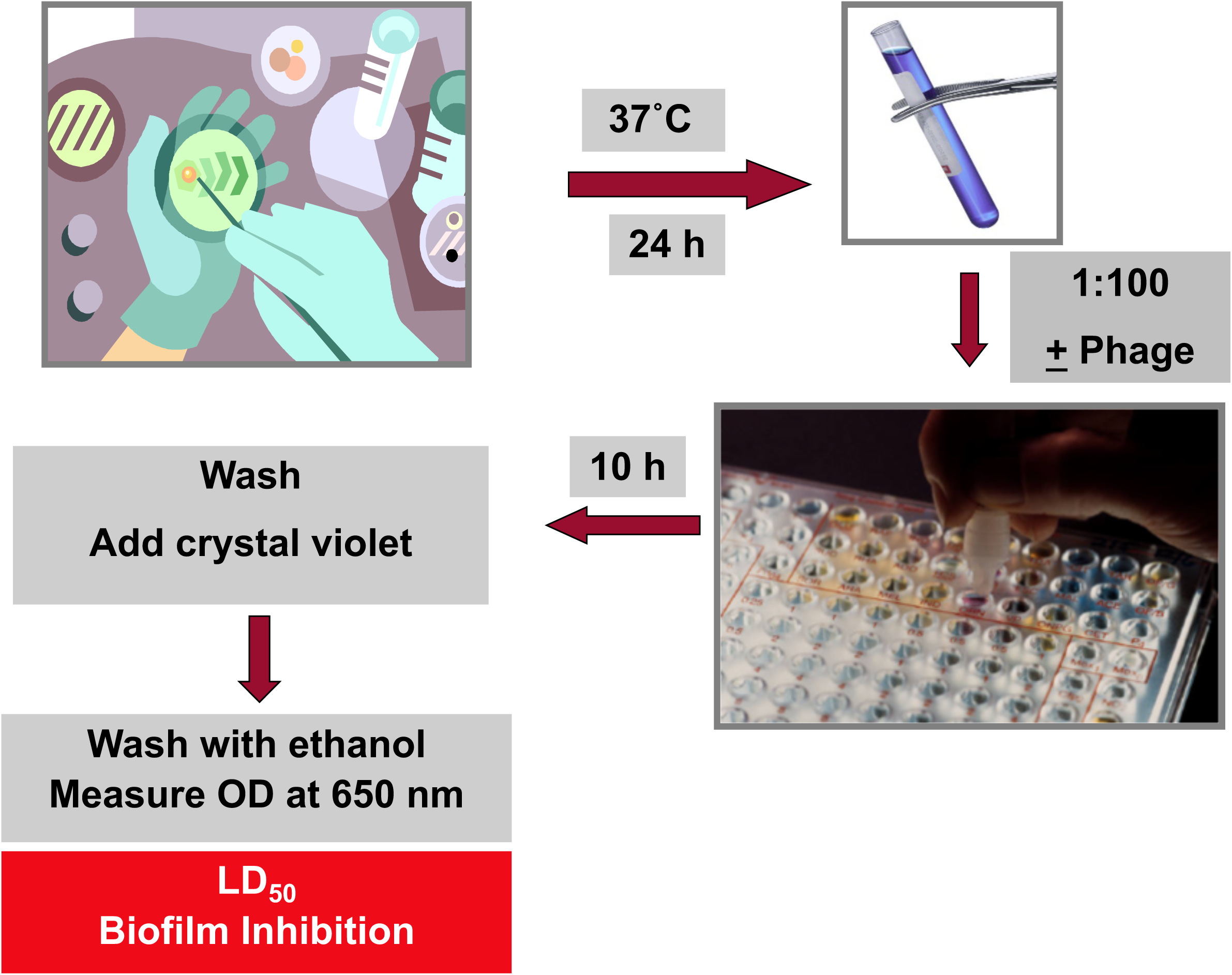
Schematic representation of the Biofilm Inhibition Assay (BIA). This assay is performed to determine if phages are useful in preventing biofilm formation. It starts with streaking the target strain on an LB agar plate. After an overnight culture a 1:100 dilution is made and inoculated in polyvinylchloride (PVC) plates along with phages at different multiplicities of infection (10, 20, 40 and 80). After an incubation period of 10 h, crystal violet is added to the plate and it is then washed with ethanol and transferred to a microtiter plate and the optical density is measured at 600 nm. The lethal dose of 50 (LD_50_) is recorded as the biofilm inhibition.

**Figure 3:**
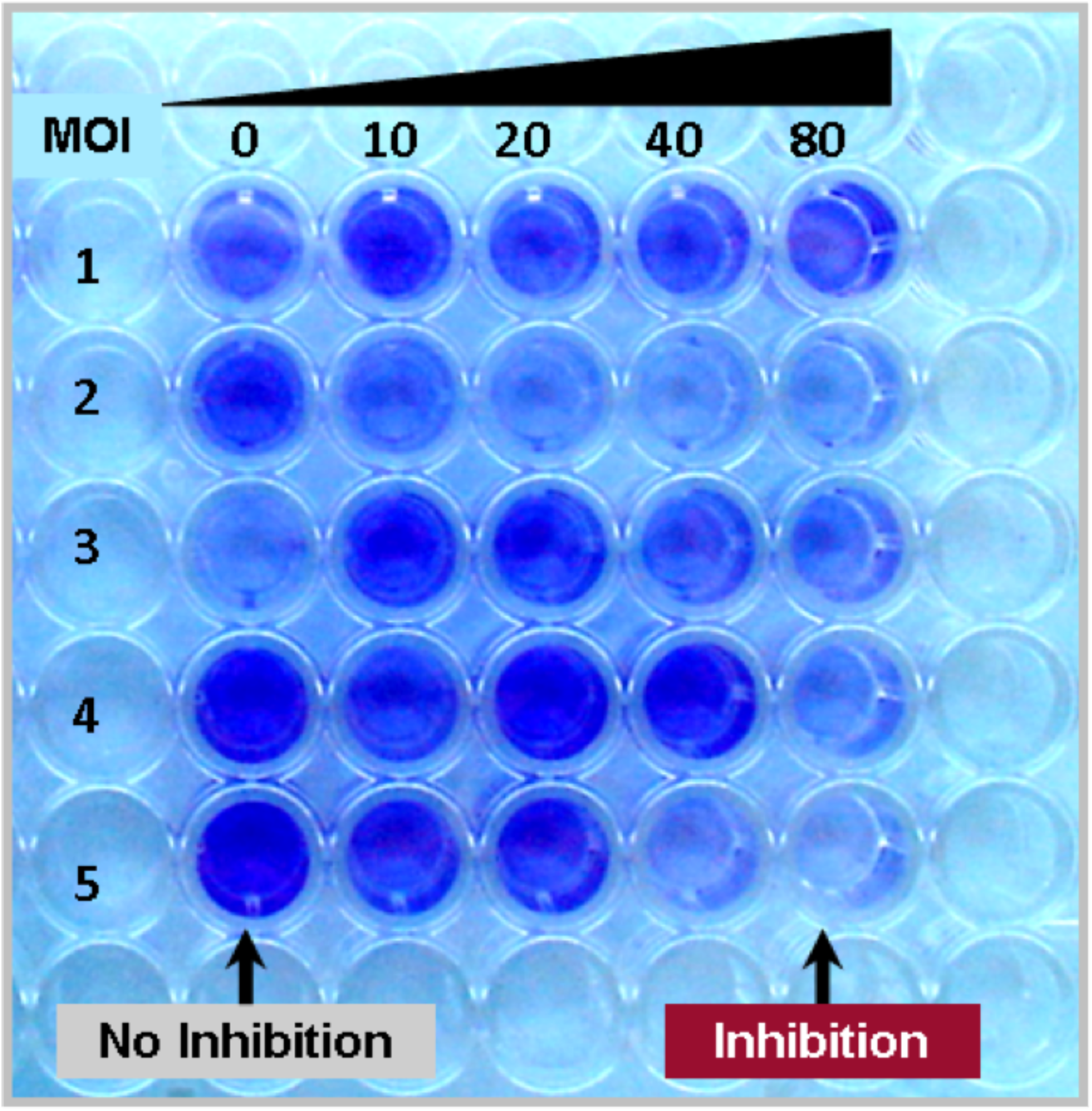
Microtiter plate in the biofilm inhibition assay: The phages were tested at MOIs of 10, 20, 40 and 80 for their ability to prevent initiation of *P. aeruginosa* biofilms. The biofilms were scored using crystal violet (CV) at an OD of 600 nm. The intensity of the color observed is proportional to the ineffectiveness of phage against biofilm initiation. In this picture, the five rows exhibit five different strains tested with phage, P140. Some of the strains required a low MOI of 10 for inhibition, eg. Row 2, whereas certain bacterial strains required higher MOIs, eg. Row 5.

To determine the amount of bound biofilm, 200 µL of 95 % ethanol was used to solubilize the CV-stained biofilm. A 125-µL aliquot of this solution was transferred to a polystyrene 96-well microtiter plate (Corning INC, Corning, NY), and the absorbance was measured at 600 nm (Packard Plate Reader Version 3.0, Ramsey, Minnesota). A higher OD reading indicated resistant bacterial strains, while a lower reading indicated more sensitive bacterial strains.

### Planktonic inhibition and biofilm eradication assay using phages

Not only it is important to inhibit the formation of biofilms, but it is also much more important to test the ability of phages to eradicate mature biofilms. To test this, four of the PCOR strains were selected based on their sensitivity to phages P140, P168 and P175B, the phages that worked best in the BIA. All these strains were inhibited at MOIs of 10 to at least two of the phages used. We included the standard prototypic strain PAO1 and its mucoid derivative PDO300 in further tests. The biofilm eradication assay was performed as described previously (90). We used the Calgary Biofilm Device (CBD), which is a two-part device. One part is a microtiter plate lid having 96 pegs, which provides the surface for uniform biofilm development. The second part is the regular microtiter 96 well plate into which the pegs fit precisely.

To perform this assay (Figure 4), the strains were inoculated in LB broth overnight at 37 °C in an air shaker. The following day, a 200 µl aliquot of a diluted (1:30) overnight culture was inoculated into the CBD wells and incubated for 24 h at 37 °C, shaking at 132 *g*. The next day the pegs were washed using 0.9 % saline solution to remove any planktonic bacterial cells. A challenge plate was prepared using a positive control (no-phage) and negative control (CAMHB broth). The selected phages, at different MOIs, were individually placed in one well of the challenge plate. This plate was incubated at 37 ^□^C in an air shaker at 320 *g* for 24 h. The next day the turbidity was determined by measuring the OD at 600 nm in a plate reader (Hewlett Packard 600, Illinois). A higher OD reading suggested the growth of surviving planktonic bacteria. An OD of less than 0.1 indicated the minimum number of phages required to inhibit planktonic bacterial growth. This was equivalent to the **m**inimum **i**nhibitory **c**oncentration (**MIC**).

**Figure 4:**
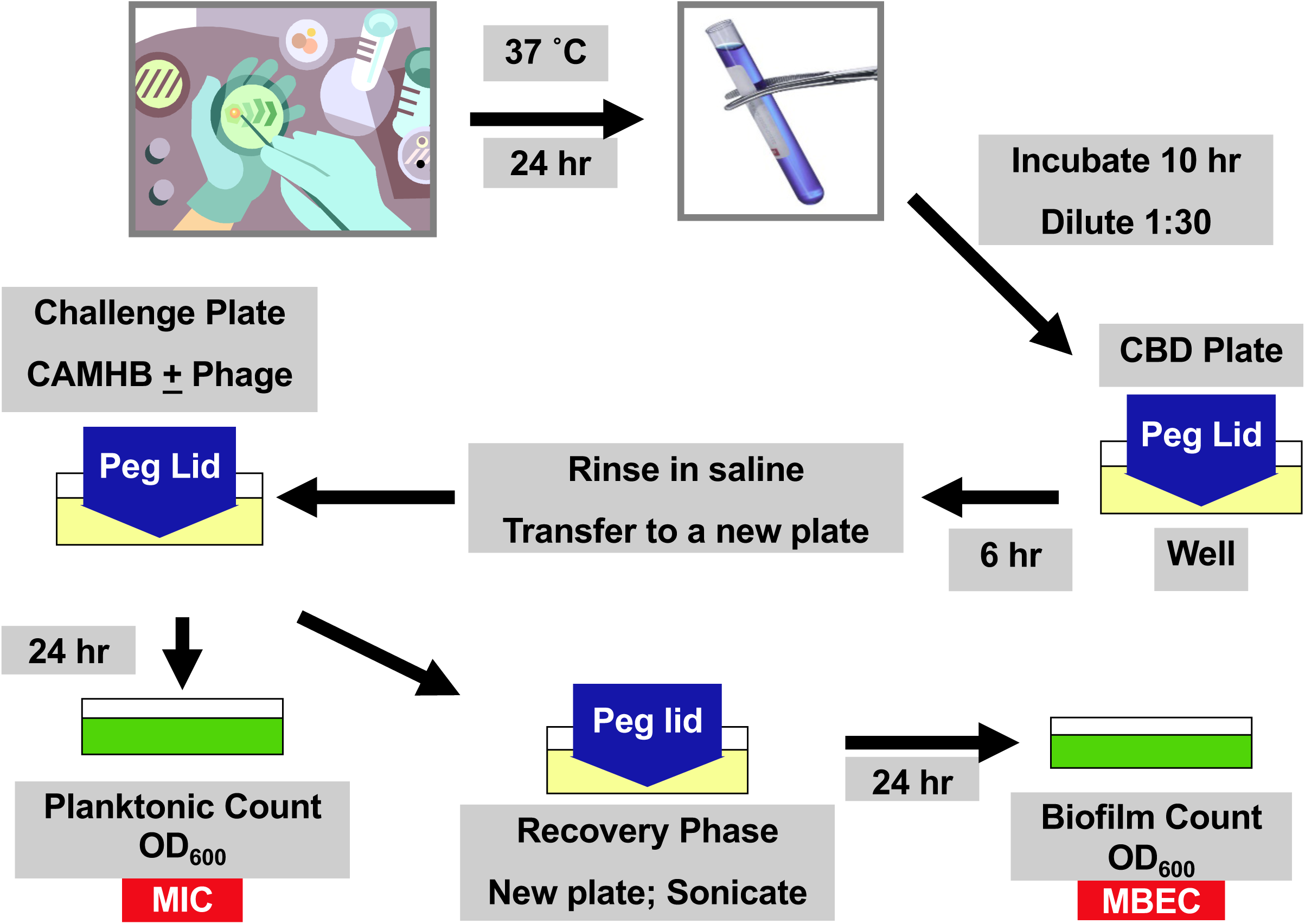
Flowchart for the biofilm eradication assay. The five-day assay begins with streaking the target strain on an LB agar plate. A day later an overnight culture is prepared which is diluted the following day 1:30 and inoculated in the CBD plate. A challenge plate is made on Day 3 with the appropriate phages and the cation-adjusted Mueller Hinton Broth (CAMHB). After an incubation period of 24 h, the (MIC) Minimum Inhibition Concentration (MIC) was measured at an OD_600_. The pegged lid is then sonicated to dislodge any surviving biofilm cells and incubated for an additional day. The minimum biofilm eradication concentration (MBEC) is measured.

The pegged lid with the coat of biofilm was rinsed in 0.9 % saline solution, placed into fresh CAMHB media and chilled on ice. The biofilm was dislodged by sonicating each peg for 10 seconds using the Microson XL Ultrasonic (Heat Systems, Inc, Farmingdale, NY). The pegged lid was discarded and the plate incubated for another 24 h to determine the number of surviving bacteria. The following day the turbidity was determined by measuring the OD at 600 nm in a plate reader. The **m**inimum **b**iofilm **e**radication **c**oncentration (**MBEC**) value was determined as the minimum concentration of phage required (when the OD is less than 0.1) for the eradication of the biofilm cells.

## RESULTS

### Screening of PCOR isolates to isolate the most effective phages

Of the 69 spatially, environmentally and geographically distinct strains of PCOR isolates tested, 48 were clinical isolates, of which 34 were from CF patients (Table 3) of which only 23 (68 %) were sensitive to phages. Out of these, 15 were sensitive at low titer (Supplement Table 1). Of the 69 strains, 23 (33 %) of them were completely resistant to all six phages. However, 46 of the 69 strains (67 %) were sensitive when tested against the six phages (Table 4).

**Table 3:** Summary of source of PCOR isolates and their sensitivity.

| Source of isolation | Total number of strains | Number of sensitive strains (%) |
| --- | --- | --- |
| CF | 34 | 23 (68) |
| Clinical | 14 | 12 (86) |
| Environmental | 12 | 8 (67) |

**Table 4:** Summary of *P. aeruginosa* PCOR strains that were screened using planktonic cells.

| Analysis | Number of strains (%) |
| --- | --- |
| Total strains screened* | 69 |
| Totals phages tested^ | 6 |
| No of sensitive strains | 46 (66.6) |
| Number of resistant strains | 23 (33.3) |
| Cocktail of phages required for 100 % sensitive strain coverage | 4 (66.6) |

\*PCOR strains: strains environmentally, spatially and geographically distinct.
^Phages used in this experiment: P105, P134, P140, P168, P175B and P182.

The most effective phage in our collection was P140, which lysed 74 % of the sensitive strains screened (Table 5). The phage that was least effective was P105, lysing only 39 % of the sensitive strains. Though P140 worked the best on the total number of strains, P134 was more effective in lysing 23 (51 %) of the sensitive strains at a low titer. Hence, at a low titer, we had 23 strains sensitive to P134, 10 strains sensitive to P140 and seven of them required P175B. Out of the other five strains that were sensitive at low titer, two required P105, two were sensitive to P182 and only one needed P168.

**Table 5:** Sensitivity of *P. aeruginosa* PCOR strains that were screened using planktonic cells.

| Phage* | Low Titer<sup>^</sup> (Unique)<sup>#</sup> | Intermediate Titer | High Titer | Total number of Strains (%)<sup>^</sup> |
| --- | --- | --- | --- | --- |
| P105 | 11 (2) | 6 | 2 | 19 (39) |
| P134 | 23 (23) | 6 | 2 | 31 (63) |
| P140 | 20 (10) | 6 (1)<sup>#</sup> | 10 | 36 (74) |
| P168 | 12 (1) | 4 | 4 | 20 (41) |
| P175B | 19 (7) | 9 | 2 | 30 (61) |
| P182 | 18 (2) | 2 | 2 | 22 (45) |

\*Phages used in this experiment: P105, P134, P140, P168, P175B and P182.
<sup>#</sup>Number of unique strains only susceptible to that phage at low titer. 45 out of the 46 strains were susceptible to one of the six phages at low titer. The 46th strain was susceptible to P140 at an intermediate titer.
<sup>^</sup>69 PCOR strains were screened against six phages for total of 414 spot titers. Low, Intermediate and High titer refers to sensitivity to $10^{-6}$ , $10^{-3}$ and $10^{-1}$ dilutions, respectively.

No single phage was observed to be effective against all sensitive strains. All six phages were needed to lyse all the strains at low or intermediate titer (Table 5). However, 100 % efficacy was reached with only four (P105, P134, P140 and P168) if higher titer was used. For example, P140 was effective against most strains but only worked against 20 bacterial strains that were sensitive to it at a low titer, i.e. even at the highest dilution of 10^−6^ and seven strains that were sensitive at a high titer, i.e. only at undiluted or 10^−1^ dilution.

### Alginate production decreases the sensitivity of the phages

The 69 strains tested included seven mucoid strains. Five (71 %) of these were resistant to all six phages. Although the sample size was small, alginate production appears to confer increased resistance to the phages tested.

### Inhibition of biofilm initiation

Since the natural mode of growth of *P. aeruginosa* is biofilms, all 46 of the sensitive strains were screened against the six phages individually in the BIA. Although all the strains’ cultures started at the same cell density at the planktonic stage, they all demonstrated different abilities to adhere to surfaces and form biofilms. This was reflected by the different optical density readings after 10 hours of incubation (Figure 5). For an example PAO1, KGN1654, KGN1648, KGN1705 had an OD of 0.658, 0.997, 1.079 and 0.684, respectively.

**Figure 5:**
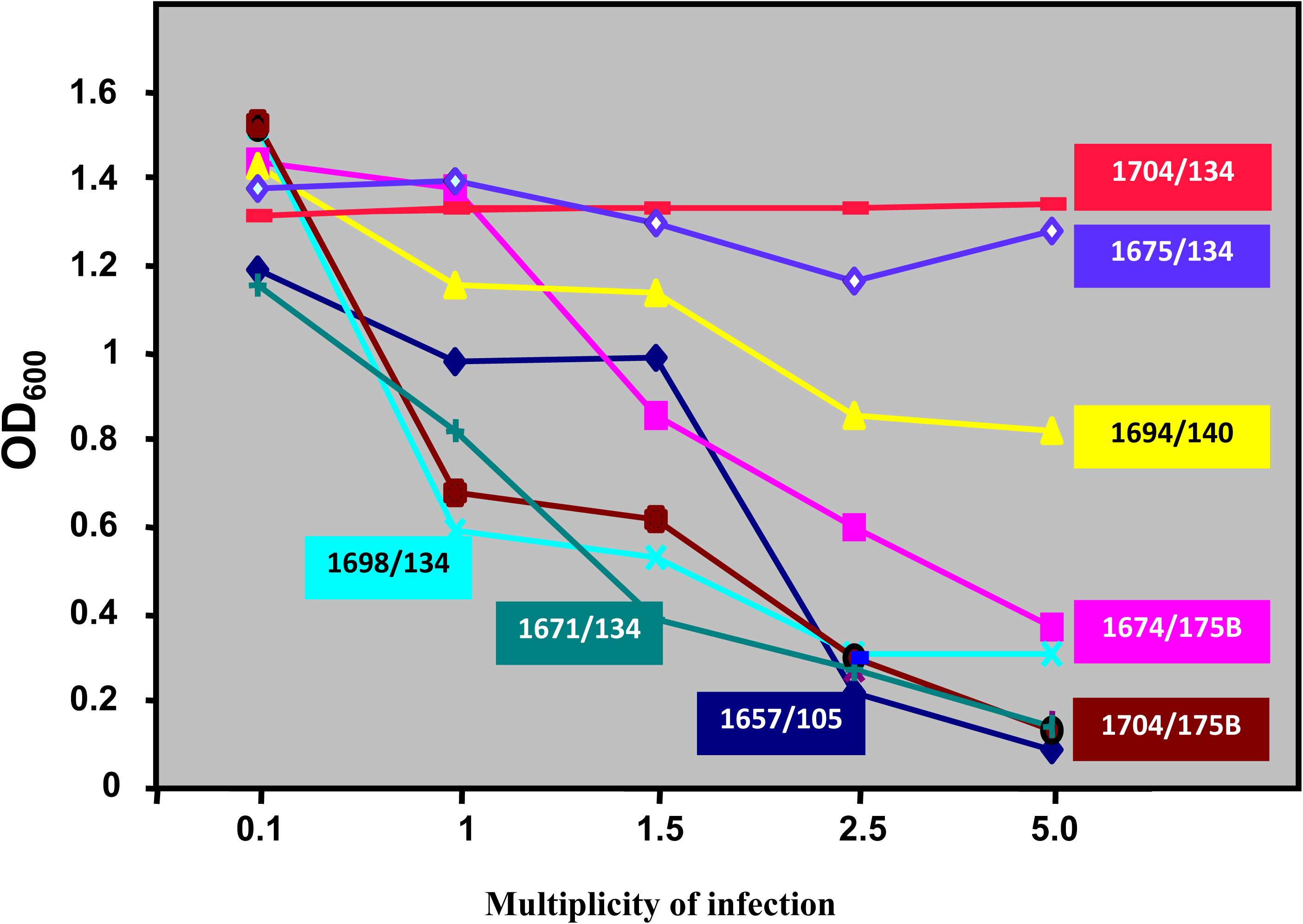
Biofilm inhibition assay with different MOIs. When lower MOIs were used some strains were sensitive, e.g. 1698, with P134 whereas some strains were resistant even at the highest MOI of 5, e.g. 1704, with P134.

Some of the 46 strains were found to be resistant at lower MOIs of 0.1, 1.0, 2.5 and 5.0 (Figure 5). However, none of the strains were completely resistant when the MOIs were increased. The LD_50_ varied from strain to strain some requiring higher MOIs as opposed to others (Figure 6). Among the strains tested, 35 of the 46 (76 %) were sensitive at the MOI of 10 (Table 6). Nine (20 %) of the bacterial strains were sensitive at an MOI of 20. When a higher MOI of 40 was tested, one more strain tested sensitive. Only two of the strains required 80, the highest MOI tested (Table 6).

**Table 6:** Sensitivity of 47 PCOR strains to different multiplicity of infections (MOIs) in the biofilm inhibition assay.

| Phage* | Multiplicity of Infection (MOI) |  |  |  | Total number of strains (%)^ |
| --- | --- | --- | --- | --- | --- |
|  | 10 | 20 | 40 | 80 |  |
| P105 | 13 (2)<sup>#</sup> | 2 | 1 | 2 | 17 (37) |
| P134 | 16 (7) | 11 (2) | 3 | 1 (1) | 31 (67) |
| P140 | 19 (19) | 9 (5) | 5 (1) | 3 (1) | 36 (78) |
| P168 | 11 (3) | 4 (1) | 7 | 0 | 21 (46) |
| P175B | 16 (3) | 6 | 6 | 2 | 30 (65) |
| P182 | 9 (1) | 8 (1) | 1 | 4 | 22 (48) |

\*Phages used in this experiment: P105, P134, P140, P168, P175B and P182.
^46 Sensitive PCOR strains were screened against six phages
<sup>#</sup>The value in parentheses refers to the number of unique strains that are only susceptible to that phage at the given MOI. 35 out of the 46 strains were susceptible to various phages at MOI of 10.

**Figure 6:**
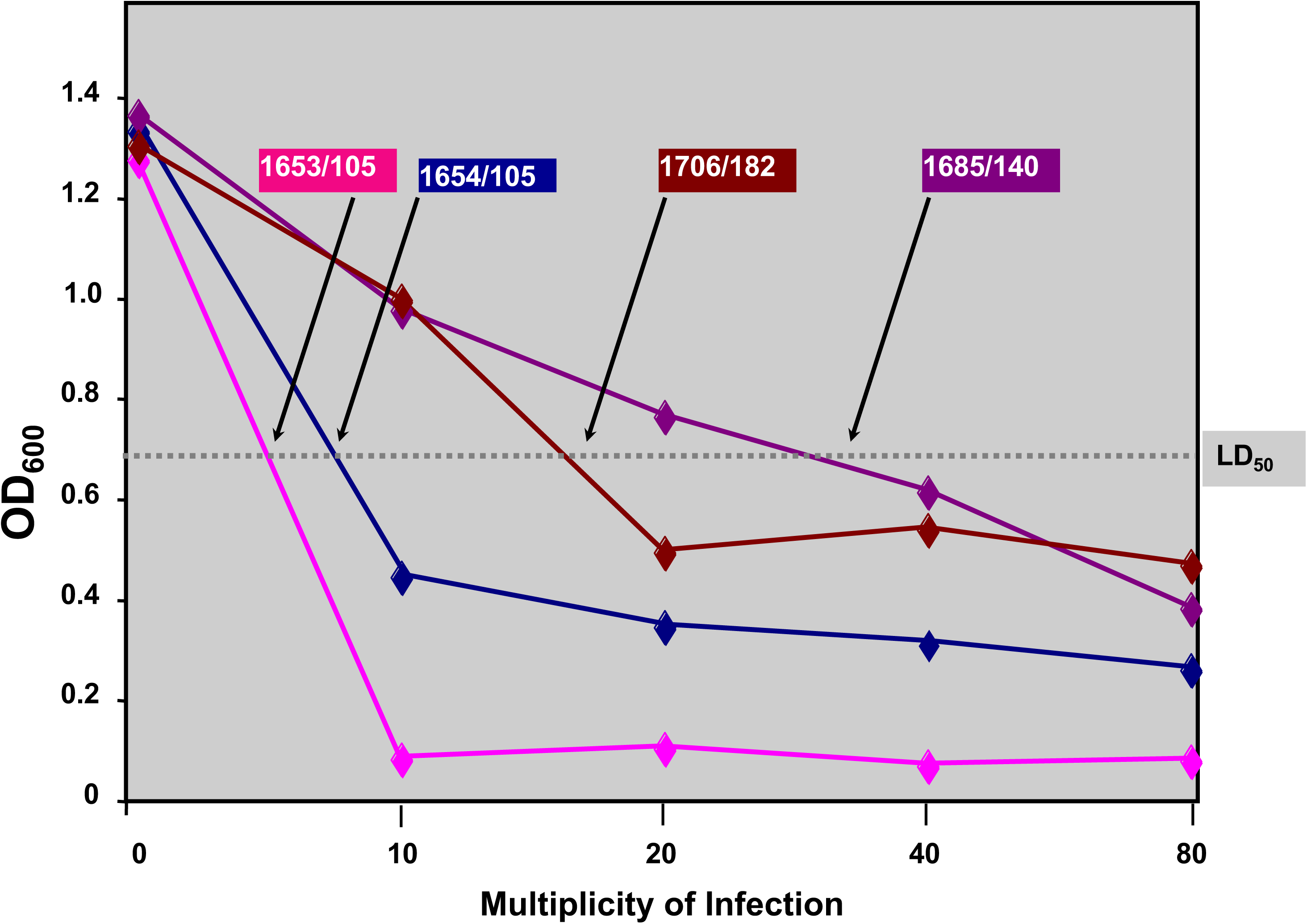
Biofilm inhibition assay showing the LD_50_ values. When the MOI was increased from 10 to 80 all the strains turned out to be sensitive. However, some required only an MOI of 10, e.g. 1653, with P105 while some required a very high MOI of 80. The LD_50_ values varied with the strains as indicated by the arrows.

The phages that were most effective against planktonic cells were not necessarily the most effective in inhibiting biofilm adhesion. For example, P134 was the most efficient phage that lysed 23 strains at low titer in the spot titer assay. However, the most effective phage against biofilm initiation was P140. The phage P140 was able to lyse 36 (78 %) of the bacterial strains and (41 %) of them were sensitive at a low MOI of 10, nine (29 %) were sensitive at an MOI of 20, while five (14 %) were sensitive at a higher MOI of 40 (Table 6).

No single phage was observed to be effective against all sensitive strains. However, for 100 % efficacy, all six phages were required (Table 6). Since it is necessary to use phages that would work at low MOIs, we had 35 (74 %) strains that were sensitive at an MOI of 10 although we had to use all six phages (Table 7). However, by increasing the MOI, we noticed that more strains could be covered with a lower number of phages (Table 7). For example, for all 47 strains to be sensitive we just needed four phages. Hence, in this case, a complete (100 %) efficiency required a cocktail of four of the six phages: P105, P140, P134 and 168. However, for 100 % efficiency much higher MOIs are required than were used in the assay for planktonic cells.

**Table 7:** Summary of the sensitivity of 47 sensitive *P. aeruginosa* PCOR strains using the Biofilm Inhibition Assay (BIA).

| MOI | Number of phages | Phages | Number of strains (%) |
| --- | --- | --- | --- |
| 10 | 6 | P105, P134, P140, P168, P175B, P182 | 35 (74) |
| 10-20 | 5 | P105, P134, P140, P168, P175B | 42 (89) |
| 10-40 | 4 | P105, P134, P140, P168 | 45 (96) |
|  | 5 | P105, P134, P140, P168, P175B | 46 (98) |
| 10-80 | 3 | P105, P134, P140 | 45 (96) |
|  | 4 | P105, P134, P140, P168 | 47 (100) |

### Dose-Dependent inhibition of biofilm formation by phages

For therapeutic purposes, it is important to establish that biofilm inhibition occurs at phage in a dose-dependent manner. The dose-dependency was tested in the BIA assay by adding phages at different MOIs (Figure 5). As controls, we also included two resistant strains, PDO300 and 1647 (data not shown). As expected, for the resistant strains, the OD measured the same at all the MOIs tested. The most sensitive strains showed the strongest inhibition at an MOI of 5 as compared to an MOI of 1 (Figure 5). For example, the strain KGN1704 tested against P175B showed a drop in the OD600 from ∼1.6 to 0.2 going from no addition of phage to MOI of 5. The dose-dependent inhibition of biofilm can also be observed at a higher MOI (Figure 6). All the sensitive strains showed this behavior (data not shown).

### Phages can be used against mature *P. aeruginosa* biofilms and prevent recolonization of planktonic cells

Out of the four strains tested three of them were sensitive to the phages. However, they required a higher MOI for both the minimum inhibition concentration (MIC) and the minimum biofilm eradication concentration (MBEC). For example, KGN1653 was sensitive at an MOI of 10 in the BIA, whereas the same strain had a MIC of 100 and MBEC of 200 (Figure 7) when tested against the same phage in the biofilm eradication assay. Similarly, the prototypic strain PAO1 needed even higher MOIs; more than 300 to eradicate the mature biofilm (Table 8).

**Table 8:**
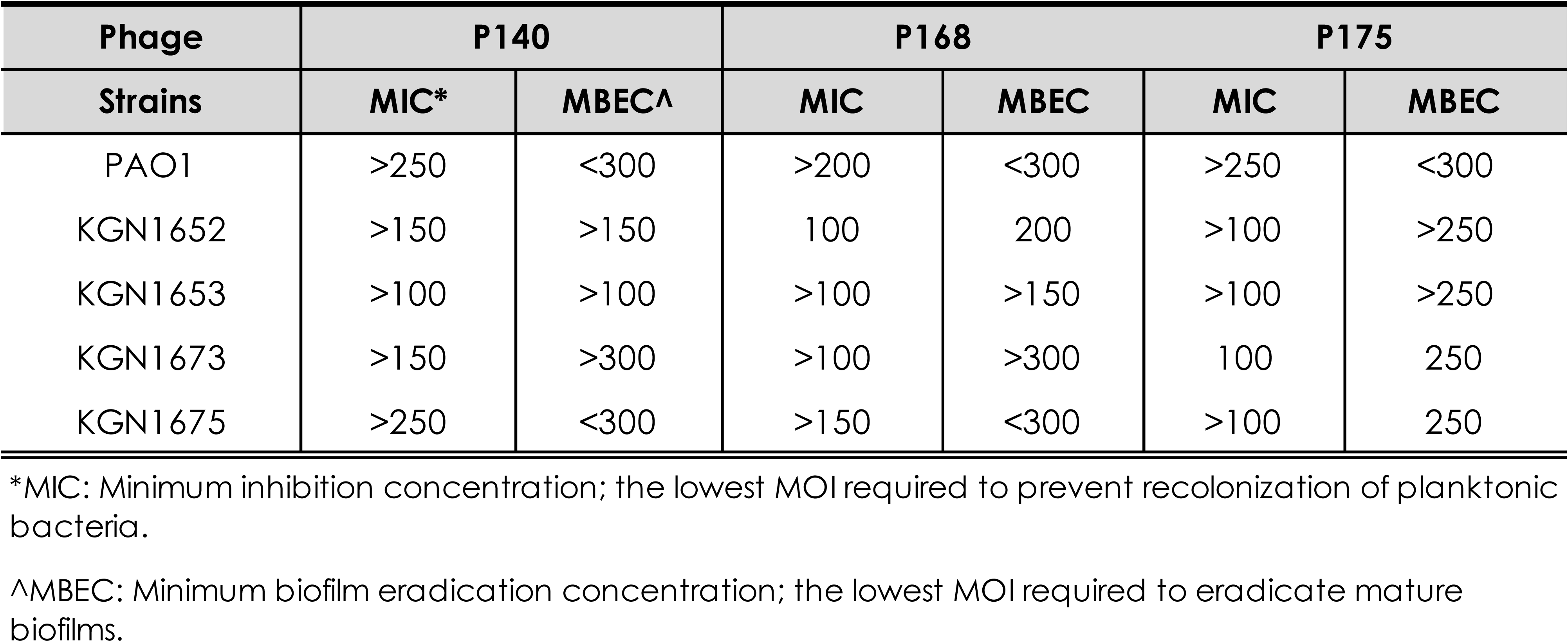
MIC and MBEC for PAO1 and the four PCOR isolates tested against P140, P168 and P175B.

**Figure 7:**
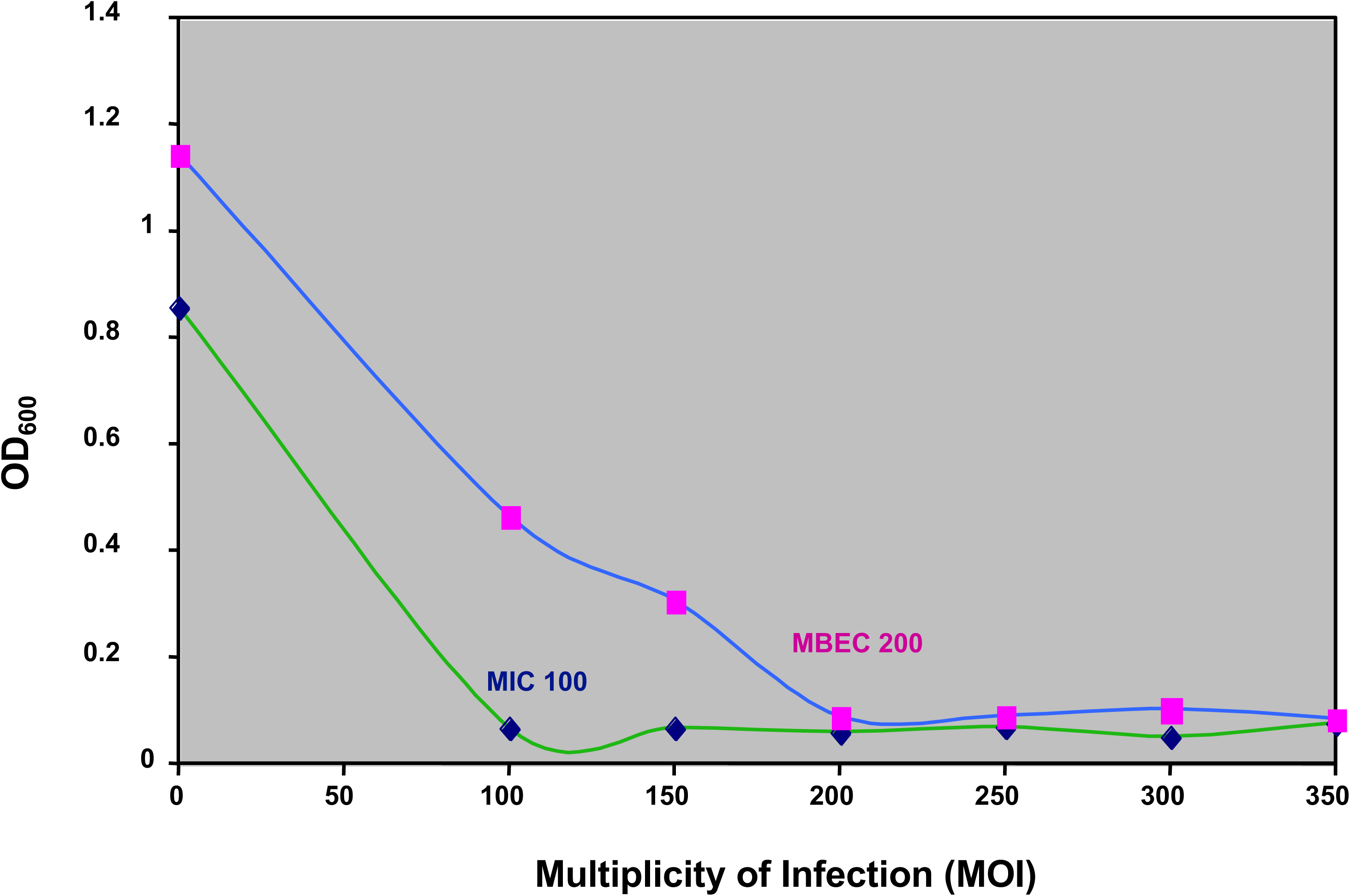
MIC and MBEC of 1653 plotted against MOI. The MOI required for MIC, that is the concentration of phage required to prevent recolonization of planktonic bacteria from the biofilms is only a 100. However, the MBEC the concentration of phage required for eradication of mature biofilm is 200.

## DISCUSSION

Experiments demonstrating the potential application of phage therapy against bacterial diseases such as diarrhea caused by *E. coli* O157, tuberculosis caused by *Mycobacterium tuberculosis* and *M. avium*, invasive gastroenteritis caused by *Vibrio vulnificus* has renewed interest in phages and their various uses (91; 92; 93; 94). *P. aeruginosa* is resistant to a large number of antibiotics (1; 2) and resistance of this bacterium has increased over time and even an increase of a thousand-fold has been observed (95). This increase in resistance raises the need to explore other effective modalities of treatment. Our results show that phages can be effective against a variety of *P. aeruginosa* strains *in vitro*. Effectiveness depends on the developmental stage of the bacterial biofilms and is dose-dependent.

### Phages are effective against a large number of *P. aeruginosa* strains

A number of clinical studies have explored the use of phages against a limited number of *P*. *aeruginosa* infections (78; 81; 82). *P. aeruginosa* strains are genomically diverse (86; 96). Ideally, a phage chosen for therapy should be effective against the diverse genome of *P. aeruginosa.* Hence, a library of PCOR isolates (Table 2) that were spatially, environmentally and geographically distinct, as well as the prototypic nonmucoid PAO1 and its isogenic mucoid variant PDO300, were tested against six phages P105, P134, P140, P168, P175B and P182. The results obtained showed that this set of phages was effective against 46 out of the 69 strains (Table 3). These results support further investigation of this method of treatment. Since 23 of the total strains tested were resistant out of which 11 were isolated from CF patients, there is a need for more phages to be isolated and tested.

No single phage was effective against all the PCOR strains. But only four of the six phages were required for 100 % efficacy (Table 7). Thus, it is very important to test the effectiveness of each of these phages against an infection before any treatment. Once a sputum culture is obtained from a patient and the bacteria isolated, it is necessary to perform a spot titer just like an antibiogram that is performed in hospital settings today. Alternatively, one could use a cocktail of phages for treatment. But before attempting this, the behavior of phage cocktails against resistant strains needs to be further elucidated. The effectiveness of a phage cocktail against *P. aeruginosa* and other pathogens has been demonstrated by using animal infection models (97). Animal studies support phage cocktails. It is believed that if cocktails are used for treatment the strains will be sensitive to some of the phages and this will be a more effective approach (98). Other studies suggest that using cocktails is important for bacterial sensitivity to phages for a longer period of time, i.e. resistance to the phages will develop much slower. Both of these factors have been well demonstrated by using *E. coli* infections in calves, piglets and lambs (73; 99) as an animal model (73) and in *P. aeruginosa* infections in human subjects (82).

Our data suggested that four of the six phages tested were required for 100 % efficacy (Table 7). Isolations of much more effective phages or construction of more virulent phages could reduce this number to maybe one or two and enhance the potential use of this therapy (100). These phages need to be effective against the bacterial strains at low titer, just like the 35 PCOR strains that were sensitive to the six phages at the highest dilution, since this will decrease the probability of any side effects. Isolation of more efficient phages can be done by using the modern understanding of bacterial virulence and targeting phages at virulence. It has been demonstrated that *E. coli* phages that were selected on the basis of superior attachment to the K1 antigen in *E*. *coli* strains were more effective against diarrhea in calves (73).

### Alginate production decreases the sensitivity of the phages

Of the *P. aeruginosa* strains isolated from the lungs of CF patients with advanced stages of disease, 85 % of the strains showed a mucoid colony phenotype (101), whereas only 1 % of the bacteria isolated from other sites of infection had a mucoid morphology (102). These observations suggest that mucoid *P. aeruginosa* cells have an added advantage, and can survive in the CF lung environment. This distinctive mucoid morphology is due to the overproduction of the exopolysaccharide alginate, an O-acetylated linear polymer of D-mannuronate and L-guluronate residues (24). This expression leads to increased resistance by *P. aeruginosa* against the host’s immune response, leading to chronic pulmonary infection and poor prognosis for the patient (103; 104). Infection with alginate-producing *P. aeruginosa* in CF patients has been associated with an overactive immune response from infection to clearance and a poor clinical condition, suggesting that alginate production is a virulence factor (23; 62; 105). This exopolysaccharide layer is known to envelop the biofilm decreasing the permeability of antibacterial drugs to the cells (27).

Five of the seven mucoid strains included in the planktonic spot titer assay were resistant to phages. This suggests that alginate production by *P. aeruginosa*, switched on due to various stress factors in the harsh lung environment of the CF patients (25), could also protect the bacteria from phage infection. Our data showed that alginate production served as a barrier against the phages. A previous study with a bacteriophage that was able to reduce the viscosity of the exopolysaccharide, thus penetrating deep within and infecting the bacteria has been reported (106). Hanlon’s study, however, used just one mucoid strain and hence this result might not apply to all *P. aeruginosa* mucoid strains.

Our study emphasizes the need to test all of the mucoid strains against more phages or, probably, a combination of phages. For example, in our preliminary tests with an additional phage, P1058, one more mucoid strain was shown to be sensitive, encouraging isolation and further study of similar phages.

### Phages can be used to prevent biofilm initiation

Biofilm formation is a natural mode of growth for *P. aeruginosa* (27); it is also an important phenotype associated with CF patients (104). The formation of a biofilm can be viewed as a developmental process having five stages: adhesion (initiation), monolayer formation, microcolony formation, maturation and dispersion (25; 28; 29). In this study, we evaluated the effectiveness of phages against biofilm initiation adhesion using the biofilm inhibition assay (BIA) (89). The phage-sensitive PCOR strains in the planktonic system were analyzed. Our results showed that each of these strains had varying adherence properties suggesting that biofilm formation is not the same with all strains. This confirmed previous findings with *P. aeruginosa* PAO1 and 31 other strains that included CF isolates (107).

As compared to the planktonic cells, the biofilm cells required higher MOI, varying from 10 to 80. None of the strains were resistant at the highest multiplicity of infection (MOI) used. This suggests that some *P. aeruginosa* strains need a higher dosage of phage for clearance. This finding is consistent with other antibacterial studies that have found an increase in resistance of the biofilm cells as compared with the planktonic form, with resistance increasing from 100 to 1000-fold (2; 90; 108; 109; 110;). These results indicate that researchers seeking phages for therapy need to isolate phages that will be effective not only on planktonic cells but also against biofilms. Since it is desirable to use smaller doses for any treatment regimen, it is necessary to identify isolated phages that are effective at low MOIs. The difference in the adhesion property and the range of MOI required to inhibit biofilm formation are reflective of *P. aeruginosa* diversity.

Our two most effective phages against biofilm initiation were P140 and P134, covering 79 % of the isolates. Therefore, a cocktail of the two effective phages might turn out to be more effective than the individual phages themselves. P134, the most effective phage on planktonic cells, covering 50 % strains at low titer, was not as effective against *P. aeruginosa* biofilms. Thus, there need not be a strong correlation between planktonic and biofilm cell sensitivity and indicates the need for future studies on biofilm susceptibility. Any phages that are isolated against *P. aeruginosa* have to be tested against both experimental systems before using them in animal or human experiments.

### Dose dependency inhibition of biofilm formation by phages

Dose-dependent studies have been conducted on *E. coli* infected with phages, and they all indicate an increase in sensitivity when challenged with higher doses of the species-specific phages (111; 112). When the biofilm inhibition assay (BIA) was performed using four different multiplicities of infections (MOIs’) (Figure 5 and 6) the lethal dose 50 (LD_50_) seemed to be dose-dependent. The higher the multiplicity of infection used, the more killing was observed.

However, higher dosages of phages can have side effects. Human subject studies in phage therapy have associated pain in the liver area reported around day 3-5; this pain, though not severe, could last for several hours (113). This might be due to the release of endotoxins, as a result of extensive lysis by the phages (79). Also in the most severe cases, fever was observed for 24 h lasting for 7-8 days (79). Treatments with high doses of phages by intravenous administration have not been recommended due to septic shock (113).

Prevention of infection for better management of CF is an important goal worth pursuing. Hence, our results suggest that the lowest MOI needed to inhibit biofilm formation and growth needs to be determined in future studies using appropriate animal models. For strains that need high MOI, we need to further isolate suitable phages or construct effective recombinant phages.

### Phages are useful in biofilm eradication

Often, treatment commences upon identification of infection. Therefore, it is important to identify phages that are effective against preformed biofilm and in preventing planktonic bacteria from recolonizing. Our study indicated that a very high MOI was required for successful eradication of biofilms. Probably, it would be more helpful to use genetically engineered phages that are more virulent by nature (114). However, our study also suggests that phages may be an important prophylactic treatment, since the bacterial strains tested were sensitive at a low MOI of 10 in the biofilm inhibition assay. Phages for prophylaxis against potential infection have been supported by many studies. For example, the study on vancomycin-resistant *Enterococcus faecium* showed that 100 % of the animals could be saved when phage was delivered within 45 min of infection, however, when phage was delivered 24 h later when the infection caused morbidity only 50 % of the animals could be saved (92). Similar studies on *E. coli* respiratory infection using broiler chickens as models, *S. aureus* infections in rabbit models and studies correlating seasonal epidemics of cholera with low phage prevalence certainly encourage phage therapy as prophylactic treatment (111; 112; 115; 116).

The increasing antibiotic resistance in this microbe has stirred interests amongst various other antibacterial communities. Using phages for treatment has great advantages and hence should be strongly considered as an alternate or as a conjunction to antibiotics. In conclusion of this study, additional phages need to be isolated from the environment, especially from CF sputum and lungs. These phages need to be tested on planktonic cells but more importantly on biofilms. Since no single phage seems to be effective on the vast and diverse genome of *P. aeruginosa* a combination of different phages needs to be considered. Further characterization of these phages might reveal a better mode of treatment. Thus, phage therapy is a promise and hope to all the unfortunate victims of this infamous pathogen.

## Acknowledgements

We thank members of the Mathee laboratory for their valuable insights, especially Alexandra Tchir for formatting the manuscript.

This work is part of an Undergraduate Honors Thesis submitted in partial fulfillment of the requirements for the degree of Bachelor of Science in Biological Sciences with Honors at the Florida International University, Miami, FL.

## Ethical Statement

Not applicable.

## Abbreviations

CF: Cystic Fibrosis
CFTR: Cystic Fibrosis Transmembrane Conductance Regulator Protein
*cftr*: cystic fibrosis transmembrane regulator gene
CV: Crystal violet
LB: Luria Bertani
OD: Optical Density
MBEC: Minimum Biofilm Eradication Concentration
*P. aeruginosa*: *Pseudomonas aeruginosa*
CBD: Calgary Biofilm Device
MIC: Minimum Inhibition Concentration
PVC: Polyvinyl Chloride
*K. pneumoniae*: *Klebsiella pneumoniae*
HIV: Human Immunodeficiency Virus
*M. avium*: *Mycobacterium avium*
MOI: Multiplicity of Infection
Phage: Bacteriophage
LD: Lethal Dose.

